# Targeting Aberrant FGFR Signaling with Infigratinib Enhances the Efficacy of BTK/PI3K Inhibitors and Bendamustine in Lymphoma Cells

**DOI:** 10.64898/2025.12.23.695986

**Authors:** Giulio Sartori, Alessandro Ghiringhelli, Eleonora Cannas, Francesca Guidetti, Alberto J. Arribas, Alex Zadro, Luciano Cascione, Chiara Falzarano, Martina Imbimbo, Matteo Moretti, Andrea Rinaldi, Emanuela Lovati, Francesco Bertoni

## Abstract

**Background:** Aberrant fibroblast growth factor receptor (FGFR) signaling sustains survival and drug tolerance in various cancers, including B-cell lymphomas. We profiled FGFR/FGF expression and evaluated the FGFR1-3 inhibitor infigratinib, both alone and in combination with standard agents, in models of mantle cell lymphoma (MCL), marginal zone lymphoma (MZL), and diffuse large B-cell lymphoma (DLBCL).

**Methods:** Twenty-eight cell lines (MCL n=10; MZL n=7, including BTK/PI3K inhibitor-resistant derivatives; ABC-DLBCL n=3; GCB-DLBCL n=8) were tested by 72-h MTT assays for single-agent and fixed-ratio combination activity. Transcriptome profiling (4, 8, 12 h) was performed in MZL Karpas1718 cells treated with infigratinib (500 nM), ibrutinib (10 nM), or the combination.

**Results:** MCL preferentially expressed FGFR3/FGF9, whereas ABC-DLBCL and MZL expressed FGFR1/FGF2; FGFR1 was further enriched in resistant MZL models, consistent with autocrine activation. Infigratinib showed dose-dependent but modest single-agent activity across the panel (median IC50 3.58 μM, indicating limited efficacy at clinically achievable exposures (≤500 nM). However, combinations were broadly synergistic: in MCL, infigratinib with ibrutinib (4/4 lines) and bendamustine (5/5) outperformed either agent alone, with weaker effects with rituximab (2/4). In MZL (parental and resistant), infigratinib synergized with ibrutinib (6/6), copanlisib (6/6), and idelalisib (4/6), restoring sensitivity in resistant derivatives; synergy occurred at infigratinib concentrations ≤500 nM. Mechanistically, RNA-seq revealed largely distinct single-agent programs: ibrutinib suppressed BCR/NF-κB/inflammatory signaling, while infigratinib preferentially repressed E2F/MYC cell-cycle modules. In contrast, the combination concomitantly downregulated NF-κB pathways and abrogated MYC target signatures, with coordinated modulation of apoptosis and DNA-repair transcripts.

**Conclusions:** FGFR1/3-driven autocrine signaling contributes to adaptive survival in MCL and MZL. Although infigratinib has limited single-agent activity, it potentiates BTK/PI3K inhibitors and bendamustine at clinically relevant concentrations and reverses acquired resistance, supporting biomarker-guided clinical evaluation of FGFR blockade in relapsed/refractory MCL and MZL.

## INTRODUCTION

The fibroblast growth factor receptor (FGFR) signaling pathway is involved in essential cellular processes, and its dysregulation has been linked to tumorigenesis and the development of several human cancers ^1-4^. Its activation is mainly ligand-dependent: binding of fibroblast growth factors (FGFs) to FGFRs leads to receptor dimerization, ultimately triggering intracellular kinase autophosphorylation cascades ^1,5^. The FGF family includes 22 members, 18 of which are secreted signaling ligands, typically divided into two main functional subfamilies based on their mode of action: the hormone-like FGFs (FGF19, 21, and 23) and the canonical FGFs (FGF1–10, 16–18, and 20)^6^. They exert their effects through four FGFRs, highly conserved single-pass transmembrane tyrosine kinase receptors ^7^. The activated FGFR triggers several downstream signaling cascades, including the RAS-MAPK-ERK, PI3K-AKT, JAK-STAT, and PLCγ-PKC pathways, which affect cell metabolism and transcription regulation, influencing cell proliferation, differentiation, and signal transduction ^1^. FGFR signaling can be activated in tumor cells through various events, including gene rearrangements, abnormal nuclear translocation, FGF ligand or FGFR overexpression, and stimuli from the tumor microenvironment ^4,8^. This pathway is aberrantly activated in 5%-10% of all human cancers, although this frequency increases to 10%-30% in urothelial carcinoma and intrahepatic cholangiocarcinoma ^2,9^. The most common genomic alteration in the FGFR family is gene amplification (66%), followed by mutations (26%), and gene rearrangements or fusions (8%) ^9^. FGFR signaling supports lymphoma-cell survival and contributes to resistance to chemotherapy and targeted agents ^2,10-16^. In DLBCL with acquired resistance to components of R-CHOP, the FGFR1/2 inhibitor fexagratinib (AZD4547) lacks single-agent activity but enhances chemotherapy and reduces the emergence of resistance ^17^. Elevated FGFR1 expression correlates with poorer outcomes in mantle cell lymphoma (MCL) ^13^. Microenvironment-derived IL-6 can trigger an FGF2–FGFR autocrine loop in lymphoma cells, which also engages EZH2, a protein positively regulated by FGFR1 ^12,13^; accordingly, FGFR inhibition induces the death of MCL cells ^12^. FGFs, including FGF2 and FGF12, have also been implicated in the pathogenesis of hairy cell leukemia ^14-16^. As a prominent oncogenic driver, FGFR has become a therapeutic target for which several targeted agents have been developed and introduced into clinical practice ^10^. Currently, three inhibitors are approved for tumors harboring FGFR alterations: erdafitinib and futibatinib for cholangiocarcinoma, and pemigatinib for urothelial carcinoma ^11,12^. Other small molecules are under development, including the FGFR1-3 inhibitor fexagratinib (AZD4547) and the multikinase inhibitors dovitinib and ponatinib ^13^. Additionally, combination trials evaluating FGFR inhibitors in conjunction with chemotherapies, immunotherapies, and antiangiogenic agents are ongoing. Infigratinib (NVP-BGJ398, BGJ398, KK8398) is a potent FGFR inhibitor with preferential specificity for FGFR1, FGFR2, and FGFR3 ^4,18^. Single-agent activity is primarily driven by genetic alterations in the FGFR pathway, including gene rearrangements, activating mutations, and amplification ^19^. Infigratinib has demonstrated promising clinical activity and a manageable adverse event profile in Phase 1-2 studies ^19-21^, leading to its FDA accelerated approval in 2021 for the treatment of cholangiocarcinoma harboring the *FGFR2* gene rearrangement. However, the confirmatory Phase 3 trial testing its efficacy against platinum-based chemotherapy was prematurely closed due to slow recruitment, leading to the withdrawal of its approval at the sponsor’s request. Analogously, the role of infigratinib as adjuvant therapy in FGFR3 aberrant high-risk resected urothelial cancer could not be established due to insufficient accrual in the phase 3 PROOF302 trial ^22,23^. The drug has also shown promising activity in children with achondroplasia ^24^, and two phase 3 trials are currently underway in this indication (NCT06164951, NCT06926491). Here, we studied FGFR and FGFs expression in lymphoma models and explored infigratinib as a single agent and in combination with established anti-lymphoma agents.

## MATERIALS AND METHODS

### Cell lines

Cell lines were obtained as described in Table S1. Cell lines were cultured under standard conditions at 37□°C in a humidified atmosphere, with 5% CO2. All media, as listed in Table S1, were supplemented with fetal bovine serum (10% or 20%) and penicillin-streptomycin-neomycin (≈5,000 units penicillin, 5 mg streptomycin, and 10 mg neomycin/mL; Sigma). Cell line identity was confirmed short tandem repeat DNA fingerprinting using the Promega GenePrint 10 System kit (B9510). All experiments were performed within 1□month of thawing, and the cells were regularly tested to be free from mycoplasma contamination using MycoAlert (Lonza).

### RNA expression datasets

RNA expression levels for cell lines and resistant models were retrieved from in-house-produced datasets (GSE221770 and GSE173984), and the status of *BCL2, MYC*, and *TP53* from a previous study ^25^.

### Immunoblotting

Cells were harvested and lysed by boiling samples in 2x Laemmli sample buffer (BioRad) supplemented with β-mercaptoethanol (Merck) for 10′. Lysates were resolved according to molecular weight by electrophoresis using Mini-PROTEAN TGX Precast gels 4–20% gradient (BioRad). After electrophoresis proteins were blotted onto nitrocellulose membrane (BioRad) by electric transfer and the membranes were blocked in TBST (20□mM Tris-HCl [pH□7.5], 150□mM NaCl, 0.1% Tween 20) with 5% nonfat dry milk (BioRad) for 1□h at room temperature (RT). The following primary antibodies were used in TBST 5% BSA buffer: FGFR1 mouse monoclonal (M19B2) (NB600-1287, Novus),Bek/FGFR2 mouse monoclonal (C-8) (sc-6930, Santa Cruz), FGFR3 mouse monoclonal (MAB766-100, ReD) and FGFR-4 mouse monoclonal (A-10) (sc-136988, Santa Cruz). Mouse monoclonal α-GAPDH (FF26A/F9, CNIO) was used in TBST with 5% nonfat dry milk. The secondary antibodies used were: ECL α-mouse IgG horseradish peroxidase-linked species-specific whole antibody (GE Healthcare). Membranes were treated with Westar ηC 2.0 chemiluminescent substrate (Cyanagen), and signals were detected using digital imaging with Fusion Solo (Vilber Lourmat).

### Treatments and proliferation assays

Infigratinib was kindly provided by Helsinn Healthcare SA. Ibrutinib, bendamustine, copanlisib, and idelalisib were purchased from Selleckchem (Houston, TX, USA). Rituximab was purchased from Roche (Basel, Switzerland). For the *in vitro* experiments, copanlisib was dissolved in 5% trifluoroacetic acid (TFA) in dimethyl sulphoxide; all other drugs were dissolved in DMSO only. Response to single drugs or drug combinations was assessed after 72 h of exposure to increasing doses of the drug, followed by an MTT [3-(4,5-dimethylthiazolyl-2)-2, 5-diphenyltetrazoliumbromide] assay. Briefly, cells were seeded in 96-well plates (non-tissue culture treated) at a density of 10,000 cells per well. Increasing doses of infigratinib diluted 1:3 (three wells for each concentration; compounds were added in a range of concentrations between 2000 and 0.33□nM). Cells were incubated for 72 hours at 37°C and 5% CO_2_. Wells containing medium only were included on each plate and served as blanks for absorbance readings. MTT (Sigma) was prepared as a 5 mg/mL stock solution in PBS and filter-sterilized. MTT solution (22□μL) was added to each well, and tissue culture plates were incubated at 37°C for 4□h. Cells were then lysed with 25% SDS lysis buffer (55□μL/well), and absorbance was read the next day at 570□nm using the Beckman Coulter-AD340. Sensitivity to single-drug treatment was evaluated by IC_50_, determined using a 4-parameter calculation upon log-scaled doses with the R software. Cell cycle distribution was evaluated on cells treated with DMSO control or infigratinib as single agents (2X IC50).

For the in vitro combination effect, a fixed ratio of infigratinib with the previously mentioned drugs was assessed. Cell viability was assessed using the MTT assay after 72 hours of compound exposure. IC50 values, isobolograms, and combination indices (CI) were estimated according to the median-effect model of Chou–Talalay ^26^ using the Synergy R package, allowing the quantitative definition for additive effect (CI□=□0.9–1.1), synergism (CI□=□0.3–0.9), strong synergism (CI□<□0.3) and antagonism/no benefit (CI□>□1.1).

### Transcriptome analysis

Initial RNA quality control was performed on the Agilent BioAnalyzer (Agilent Technologies, California, USA) using the RNA 6000 Nano kit (Agilent Technologies) and concentration was determined with the Invitrogen Qubit (Thermo Fisher Scientific) using RNA BR reagents (Thermo Fisher Scientific). Total RNA samples were prepared for RNA-Seq with the NEBNext rRNA Depletion kit, the NEBNext Ultra Directional RNA Library Prep Kit for Illumina and NEBNext Multiplex Oligos for Illumina (New England BioLabs Inc.). Sequencing was performed using a NextSeq 500 with the NextSeq 500/550 High Output Kit v2 (150□cycles PE; Illumina). All data will be available at the National Center for Biotechnology Information (NCBI) Gene Expression Omnibus (GEO) (http://www.ncbi.nlm.nih.gov/geo) database.

### Data mining

For RNA-Seq data, we evaluated the reads quality with FastQC (v0.11.5) ^27^ and removed low-quality reads/bases and adaptor sequences using Trimmomatic (v0.35) ^28^. The trimmed-high-quality sequencing reads were aligned using STAR ^29^ to the reference genome (HG38). The HTSeq-count software package ^30^ was then used to quantify gene expression with GENCODE v22 as gene annotation. We sub-setted the data to genes with a counts-per-million value greater than five in at least one sample. The data were normalized using the ‘TMM’ method from the edgeR package ^31^ and transformed to log2 counts-per-million using the edgeR function ‘cpm.’ The differential gene expression of each comparison of interest was computed using limma based on TMM-normalization and voom transformation ^32^. Functional annotation was done with *g:Profiler* using gene sets from the Molecular Signatures Database (MSigDB v5.1) [33] (Hallmark, c2.all, c5.bp, c6), SignatureDB [34] and gene sets obtained from different publications as reported. *Standard settings were used for g:Profiler* data mining [35]. *Signatures with absolute log fold change*□*>*□*0*.*1* and *adj*.*P*□*<*□*0*.*05* were considered as biologically relevant. Functional analysis was also performed using GSEA (Gene Set Enrichment Analysis) with the MSigDB (Molecular Signatures Database) C2-C7 gene sets (18, 19) ^33^, and SignatureDB database ^34^. Statistical tests were performed using the R environment (R Studio console; RStudio, Boston, MA, USA).

## RESULTS

### Lymphoma models derived from NF-κB-driven lymphoma exhibit active FGF signaling, sustained through autocrine loops

We first analyzed FGFR1–4 and FGF ligands expression across a panel of B-cell lymphoma cell lines using in-house RNA-Seq data ^35^ (Supplementary Figure S1A). MCL cells displayed high levels of FGF9 together with its cognate receptors FGFR1 and FGFR3 (Figure 1A). FGFR1 and its ligand FGF2 were strongly expressed in activated B-cell-like (ABC) DLBCL and marginal zone lymphoma (MZL) models. These findings suggest that MCL, ABC-DLBCL, and MZL maintain an active, autocrine FGFR signaling. Notably, all three subtypes are also characterized by constitutive NF-κB activity ^36^. Immunoblotting was used to confirm FGFR1/3 expression at the protein level and were also included a series of models of secondary resistance to BTK and PI3K inhibitors derived from the MZL VL51 and Karpas1718 cell lines ^37-40^. The ABC DLBCL cell lines OCI-Ly3 and U2932 showed higher FGFR1 expression than the MCL (SP53, Z138, JEKO1) and germinal center B-cell type (GCB) DLBCL (VAL and OCI-Ly1) models (Figure 1B and 1C; Supplementary Figure S1B and S1C). Conversely, FGFR3 protein levels were higher in MCL cell lines than in MZL and GCB DLBCL models (Supplementary Figure S1B and S1C). FGFR1 protein expression was also highly expressed in the MZL models derived from cells with acquired resistance to BTK/PI3K inhibitors (Figure 1B and 1C; Supplementary Figure S1B and S1C). FGFR2 and FGFR4 showed comparable expression across the analyzed samples (Supplementary Figure S1 and S1C). Two multiple myeloma cell lines (U266 and JJN3) were used as positive controls, and they expressed FGRF1 or FGFR3 protein levels like those of Karpas1718-IDE and MCL cells, respectively (Supplementary Figure S1B and S1C). Overall, RNA and protein expression data suggest that MCL, ABC DLBCL, and MZL cells exhibit autocrine FGFR activity, with FGFR1 being more relevant in ABC DLBCL and MZL, and FGFR3 in MCL.

**Figure 1.**
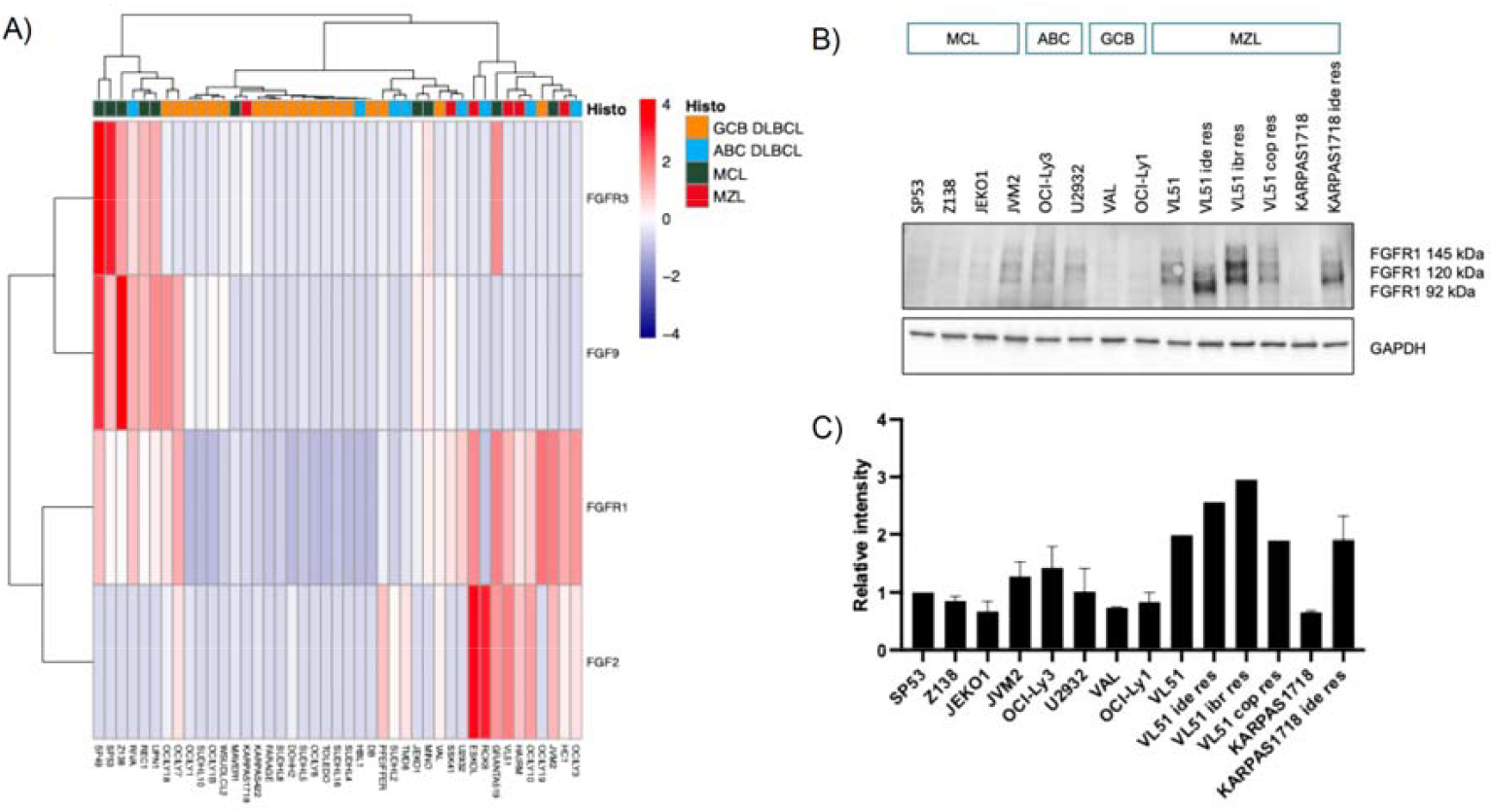
FGFR expression in lymphoma cell lines: **A)** Heatmap showing expression of FGFR1/3 and FGF2/9 across a panel of cell lines derived from MCL, ABC-DLBCL, and germinal center B-cell (GCB) DLBCL; heatmap is based on log2 CPM (counts per million). The scale ranges from low (-4, blue) to high (+4, red). Each horizontal represents the values for one cell line. **B)** Immunoblot for FGFR1 protein in a subset of selected cell lines and **C)** its protein quantification expressed as a ratio to SP53; data is the average with the second replicate shown in Supplementary Figure 1.

### Infigratinib exerts limited single-agent anti-tumor activity in B-cell lymphoma models

Since some lymphoma models showed an active FGFR signaling, we assessed the anti-tumor activity of the FGFR inhibitor infigratinib across 28 B-cell lymphoma cell lines, derived from MCL (n. = 10), MZL (n. = 7, including the resistance models), ABC DBCL (n. = 3), and GCB DLBCL (n. = 8). After 72 hours of exposure, infigratinib induced a dose-dependent anti-proliferative effect in all models, with a median IC50 of 3.58 μM (Figure 2; Supplementary Figure 2; Supplementary Table 1). The most sensitive lines were OCI-LY7 (IC50, 0.4 μM) and Karpas1718-IDE (IC50, 0.59 μM), which showed enhanced sensitivity compared with their parental cells (IC50, 2.31 μM). Similarly, the models with resistance to BTK inhibitors, derived from prolonged exposure to the BTK inhibitor ibrutinib (VL51-IBRU; 3.8 μM) or the PI3Kδ Inhibitor idelalisib (VL51-IDE; 5.3 μM), were more sensitive than the parental cells (8.3 μM). Besides MZL cell lines and their resistant derivatives, there was no correlation between sensitivity to infigratinib and FGFR1/FGFR3 expression. We then performed cell cycle analysis on a panel of lymphoma cell lines selected based on their sensitivity to infigratinib, including Karpas1718 and its Karpas1718-IDE derivative with acquired resistance to BTK and PI3K inhibitors, four GCB DBCL (OCI-LY-7, PFEIFFER, FARAGE, and VAL) and two ABC DLBCL (TMD8 and U2932) cell lines. Treatment with infigratinib at 2-times the IC50 reduced proliferative fractions and increased sub-G0 populations in these sensitive models (Supplementary Figure 3). These findings indicate that infigratinib impairs the proliferation and survival of B-cell lymphoma cell lines but, taking into consideration a 500 nM threshold, corresponding to the clinically achievable concentration ^19,41^, its efficacy as a single agent must be considered limited.

**Figure 2.**
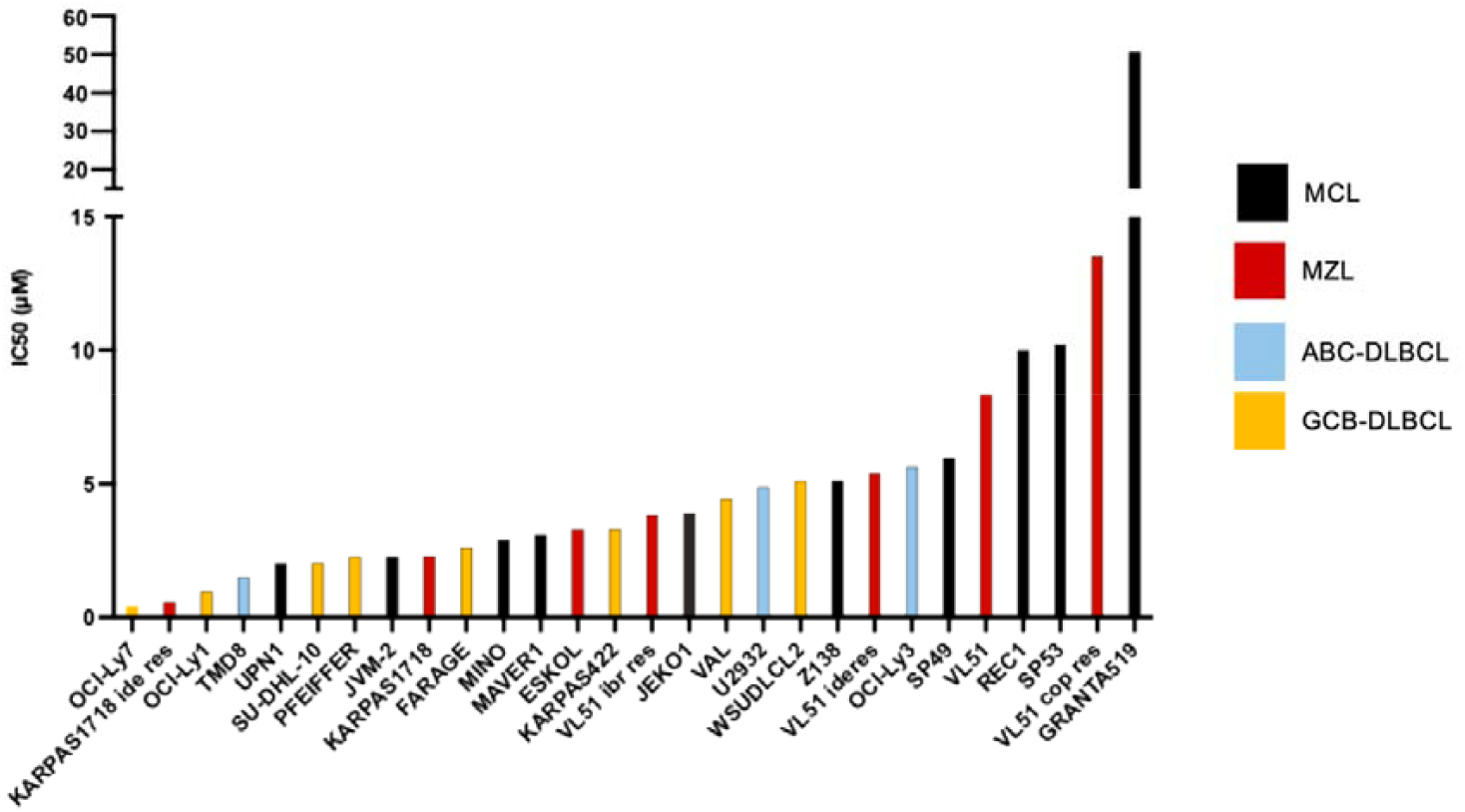
Infigratinib IC50 values in lymphoma cell lines: **A)** Barplot showing IC50 (μM) for each cell lines divided in MCL (10), MZL (7), ABC (3) and GCB (8) DLBCL.

### Infigratinib enhances the activity of targeted and cytotoxic agents in MCL and MZL models

Lymphoma cells expressed FGFR receptors and ligands, and despite the limited single-agent activity, we tested infigratinib in combination with standard therapies, hypothesizing that inhibiting the signaling could enhance the effectiveness of anti-lymphoma agents. We combined infigratinib with the BTK inhibitor ibrutinib, the chemotherapeutic agent bendamustine, and the anti-CD20 monoclonal antibody rituximab in four MCL cell lines (JEKO1, REC1, SP49, and Z138). Infigratinib synergized with ibrutinib (4/4) and bendamustine (4/4), but only weakly with rituximab (2/4) (Figure 3, Supplementary Table S2). We also explored the benefit of adding the FGFR inhibitor to targeted agents in the MZL models, including the models of secondary resistance (Karpas1718, Karpas1718-IDE, VL51, VL51-IDE, VL51-IBRU, VL51-COP). The infigratinib/ibrutinib combination was synergistic in VL51 and Karpas1718 parental cells and in their derivative resistant cells (6 out of 6 models) (Figure 3, Supplementary Table S2). The combination was more effective in the models derived from Karpas1718 than from VL51, consistent with the sensitivity to infigratinib as a single agent. Similarly, infigratinib synergized with the PI3Kα/δ inhibitor copanlisib and the PI3Kδ inhibitor idelalisib in 6/6 and 4/6 MZL models (Figure 3, Supplementary Table S2), restoring the sensitivity to the targeted agents in the resistant cells. These findings showed that infigratinib improves the efficacy of bendamustine, BTK inhibitors, and PI3K inhibitors, also at concentrations lower than 500 nM, a clinically achievable concentration ^19,41^, highlighting its potential as part of a combination strategy to overcome resistance and enhance therapeutic response.

**Figure 3.**
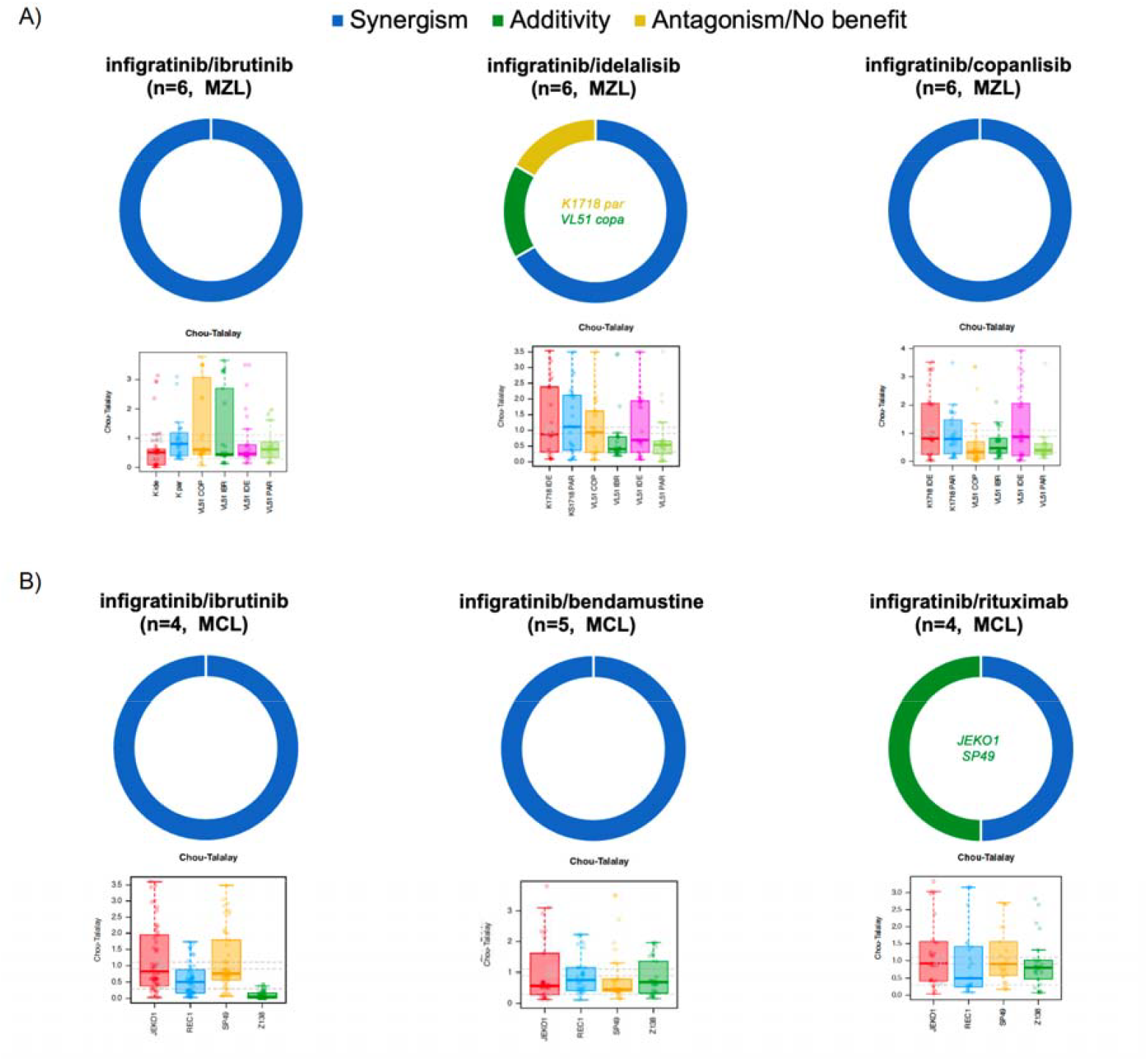
Combination activity of infigratinib in MZL models with acquired resistance and MCL models. **A)** In MCL cell lines, infigratinib was combined with ibrutinib (BTK inhibitor), bendamustine (chemotherapy), or rituximab (anti-CD20 monoclonal antibody). **B)** In MZL models (Karpas1718 and VL51) of secondary resistance to FDA-approved small molecules (ibrutinib, idelalisib, and copanlisib), infigratinib was combined with ibrutinib (BTK inhibitor) or idelalisib (PI3K δ inhibitor), or copanlisib (pan-class PI3Kδ/α inhibitor).

### Complementary mechanisms of action underlie the synergistic effect of infigratinib and ibrutinib

We then focused on the combination of infigratinib and ibrutinib, due to established use of BTK inhibitors in the clinical setting. To explore the mechanisms sustaining the observed synergy, we performed transcriptome profiling in the MZL Karpas1718 cell line treated with infigratinib or ibrutinib at clinically achievable concentrations (500 nM and 10 nM, respectively), or the combination of both agents. DMSO-treated cells were used as controls and RNA was extracted after 4, 8, and 12 hours of exposure. In line with its limited anti-tumor effect as a single agent, infigratinib alone had a modest impact on global transcription compared to ibrutinib (Figure 4A, Supplementary Table S3), with largely distinct expression profiles (Figure 4B, Supplementary Table S3). Ibrutinib suppressed genes involved in the B cell receptor (BCR) signalling (TNF and NF-κB targets) and in inflammatory response (cytokine-mediated and JAK-STAT targets), while infigratinib upregulated these pathways (Supplementary Figure 4 and Supplementary Table S3). When the two drugs were combined, a downregulation of these pathways was observed, with several key transcripts being modulated, including TNF, NFKB1/2, RELA/B, BTK, CD40, EGR1/2, CCL3/4, DDX21, CXCR4 and IRAK4 (Supplementary Table S3). Conversely, genes involved in cell cycle (including E2F, PLK1, Aurora kinase B targets) and MYC-driven pathways were repressed by infigratinib but upregulated by ibrutinib (Supplementary Figure 4 and Supplementary Table S3); combined treatment strongly suppressed these genes, including E2Fs, CDKs, PLK1 and AURKA. Notably, MYC signaling (target genes) was completely abrogated only under combination treatment, with a greater extent compared to infigratinib alone (Figure 4C, Supplementary Table S3). In addition, the combination upregulated genes involved in apoptosis, Wnt/β-catenin, and DNA repair programs, with modulation of TP53 targets (TP53BP2, TP53INP1), ATM, BRCA1/2, and WNT ligands (Supplementary Table S3). Finally comparing the genes modulated by infigratinib with publicly available data for other drugs (L100038 ^42^ and GDSC39 ^43^), we found an overlap with Src and PI3K/mTOR inhibitors, underscoring complementary pathway inhibition (Supplementary Figure 5). Overall, these data indicate that infigratinib and ibrutinib exert different transcriptional effects that converge, when combined, in a more profound suppression of oncogenic signaling and cell cycle programs, thereby providing a mechanistic basis for their synergistic anti-tumor activity.

**Figure 4.**
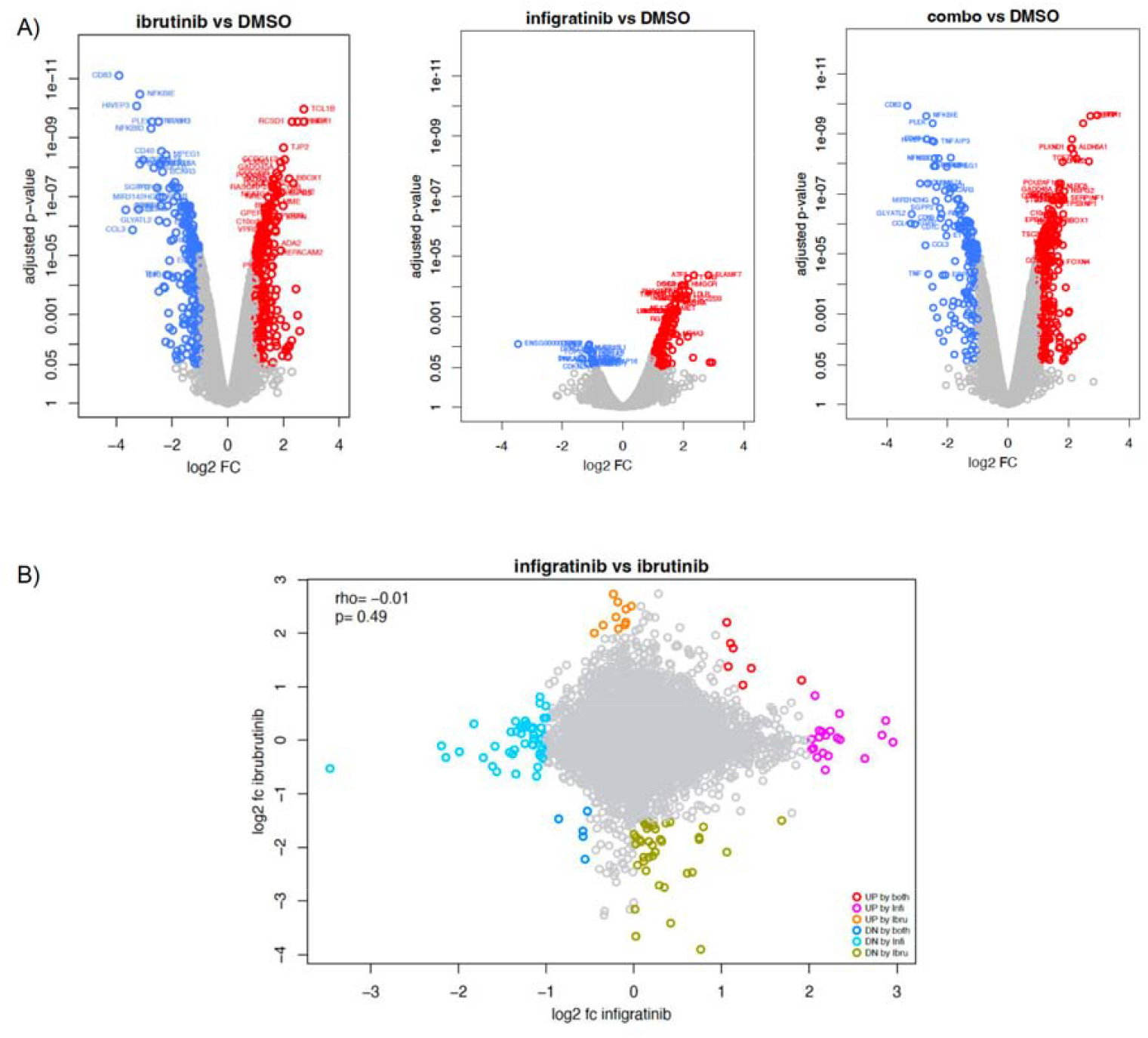
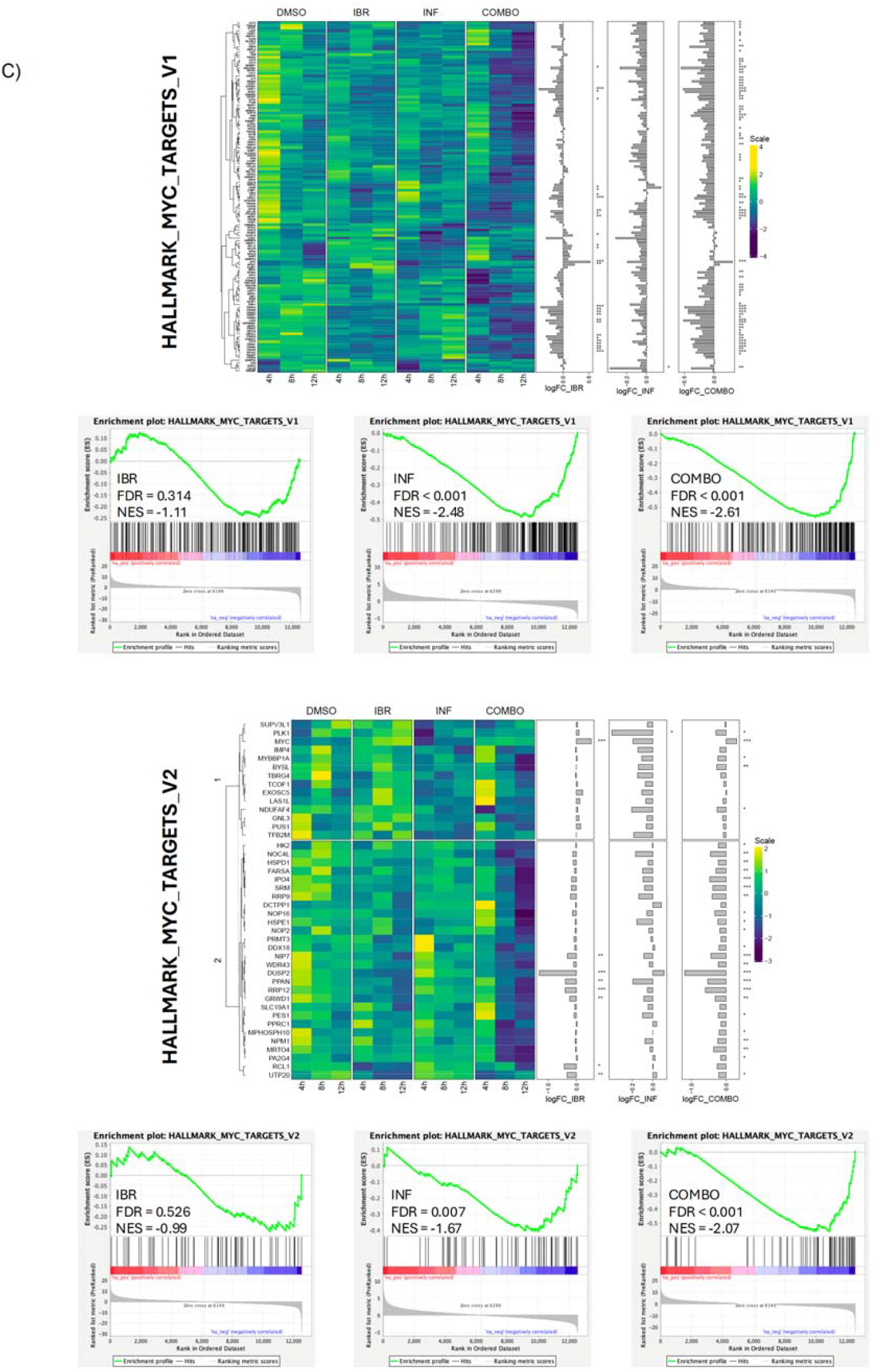
Transcriptome profiling in the MZL model KARPAS1718 cells exposed to infigratinib and ibrutinib or to the combination of the two. A) Volcano plots of ibrutinib vs DMSO (left), infigratinib vs DMSO (middle), or combination vs DMSO (right), up-regulated genes in red and down-regulated in blue. B) Correlation plot between infigratinib and ibrutinib log2 fold change. C) Differentially enriched MYC Hallmark genesets (MSigDB, Broad Institute) across treatments. FDR for false discovery rate, NES for normalized enrichment score.

## DISCUSSION

Here, we showed that MCL, ABC DLBCL, and MZL cells exhibited autocrine FGFR signaling, with FGFR1 and FGFR3 serving as dominant drivers, depending on the subtype. Indeed, FGFR1 was strongly expressed in MZL, ABC-DLBCL, and MCL, while FGFR3 was primarily expressed in MCL. Importantly, FGFR1 expression was particularly enriched in resistant MZL models, suggesting a role in therapy escape. The pharmacological inhibition of the FGFR axis with infigratinib as a single agent induced modest dose-dependent anti-tumor activity across B-cell lymphoma models, when compared with FGFR-addicted solid tumors ^19,20^. Nonetheless, we observed a clear therapeutic advantage when the FGFR inhibitor was combined with standard treatments, supporting the notion that FGFR signaling contributes to adaptive resistance mechanisms in lymphoma. Notably, the addition of infigratinib to either BTK inhibition or bendamustine enhanced efficacy across all MZL and MCL models tested. Moreover, in models with acquired resistance to BTK or PI3K inhibitors, infigratinib partially restored sensitivity to BCR pathway inhibition. Importantly, synergistic effects were achieved at clinically relevant concentrations of infigratinib (<500 nM), indicating that activation of the FGFR axis represents a compensatory survival mechanism exploited by lymphoma cells expressing FGFRs and their ligands to circumvent BCR blockade. These findings are well supported by the literature ^10,12,13,17^, which also highlights that, in patients, the effect may be more pronounced due to the production by TME cells of FGFs and cytokines that have a positive impact on FGFR signaling.

The strongest and most clinically relevant effect was observed in combination with BTK inhibition. Mechanistically, the effects of the BTK inhibitor and the FGFR inhibitor converged on complementary

pathways: ibrutinib suppressed BCR/NF-κB signaling, while infigratinib inhibited MYC- and E2F-driven proliferation. Their combination resulted in stronger repression of MYC programs and robust modulation of survival transcripts such as EGR1/2, IRAK4, and BIRC3 and pathways such as NF-κB and cell cycle, leading to an increased cytotoxic activity of ibrutinib. In conclusion, from a translational standpoint, based on the data we presented, FGFR inhibition seems worth of being explored in combination with BTK inhibitors or bendamustine in patients with relapsed/refractory MCL and MZL. Future studies should focus on biomarker-driven patient selection, prioritizing FGFR1/3 expression and resistance phenotypes.

## Declarations

### Authors’ contributions

GS performed experiments, analyzed and interpreted data, performed data mining, prepared the figures, and co-wrote the manuscript.

AG, EC, FG, AJA, AZ, and CF performed experiments.

LC performed data mining.

AR performed transcriptome profiling.

MI: co-wrote the manuscript.

MM: provided resources.

EL: provided resources and interpreted data.

FB: designed research, interpreted data, and co-wrote the manuscript.

All authors reviewed and accepted the final version of the manuscript.

## References

1. Zheng J, Zhang W, Li L, et al. Signaling Pathway and Small-Molecule Drug Discovery of FGFR: A Comprehensive Review. Front Chem. 2022;10:860985.

2. Brooks AN, Kilgour E, Smith PD. Molecular Pathways: Fibroblast Growth Factor Signaling: A New Therapeutic Opportunity in Cancer. Clinical Cancer Research. 2012;18(7):1855–1862.

3. Xie Y, Su N, Yang J, et al. FGF/FGFR signaling in health and disease. Signal Transduction and Targeted Therapy. 2020;5(1):181.

4. Katoh M. FGFR inhibitors: Effects on cancer cells, tumor microenvironment and whole-body homeostasis (Review). Int J Mol Med. 2016;38(1):3–15.

5. Chae YK, Ranganath K, Hammerman PS, et al. Inhibition of the fibroblast growth factor receptor (FGFR) pathway: the current landscape and barriers to clinical application. Oncotarget. 2017;8(9):16052–16074.

6. Beenken A, Mohammadi M. The FGF family: biology, pathophysiology and therapy. Nat Rev Drug Discov. 2009;8(3):235–253.

7. Eswarakumar VP, Lax I, Schlessinger J. Cellular signaling by fibroblast growth factor receptors. Cytokine Growth Factor Rev. 2005;16(2):139–149.

8. Bayle A, Martin-Romano P, Loriot Y. FIGHT against FGF/FGFR alterations: what are the next steps? Ann Oncol. 2022;33(5):460–462.

9. Helsten T, Elkin S, Arthur E, Tomson BN, Carter J, Kurzrock R. The FGFR Landscape in Cancer: Analysis of 4,853 Tumors by Next-Generation Sequencing. Clin Cancer Res. 2016;22(1):259–267.

10. Sacco A, Federico C, Giacomini A, et al. Halting the FGF/FGFR axis leads to antitumor activity in Waldenström macroglobulinemia by silencing MYD88. Blood. 2021;137(18):2495–2508.

11. Giacomini A, Taranto S, Gazzaroli G, et al. The FGF/FGFR/c-Myc axis as a promising therapeutic target in multiple myeloma. J Exp Clin Cancer Res. 2024;43(1):294.

12. Sehgal L, Jain N, Khashab T, wang x, Neelapu S, Samaniego F. Abstract 3286: Targeting mir101/ EZH2/NFkb axis by FGFR1 inhibitor in mantle cell lymphoma. Cancer Research. 2016;76(14 Supplement):3286–3286.

13. Sircar A, Singh S, Xu-Monette ZY, et al. Exploiting the fibroblast growth factor receptor-1 vulnerability to therapeutically restrict the MYC-EZH2-CDKN1C axis-driven proliferation in Mantle cell lymphoma. Leukemia. 2023;37(10):2094–2106.

14. Aziz KA, Till KJ, Chen H, et al. The role of autocrine FGF-2 in the distinctive bone marrow fibrosis of hairy-cell leukemia (HCL). Blood. 2003;102(3):1051–1056.

15. Forconi F, Rinaldi A, Kwee I, et al. Genome-wide DNA profiling identifies an unstable profile with recurrent imbalances predicting outcome in chronic lymphocytic leukemia with 17p deletion. Br J Haematol. 2008;143(4):532–536.

16. Arribas AJ, Napoli S, Gaudio E, et al. Secreted Factors Determine Resistance to Idelalisib in Marginal Zone Lymphoma Models of Resistance. Blood. 2019;134(Supplement_1):2569–2569.

17. Chen J, Ge X, Zhang W, et al. PI3K/AKT inhibition reverses R-CHOP resistance by destabilizing SOX2 in diffuse large B cell lymphoma. Theranostics. 2020;10(7):3151–3163.

18. Guagnano V, Furet P, Spanka C, et al. Discovery of 3-(2,6-dichloro-3,5-dimethoxy-phenyl)-1-{6-[4-(4-ethyl-piperazin-1-yl)-phenylamino]-pyrimidin-4-yl}-1-methyl-urea (NVP-BGJ398), a potent and selective inhibitor of the fibroblast growth factor receptor family of receptor tyrosine kinase. J Med Chem. 2011;54(20):7066–7083.

19. Guagnano V, Kauffmann A, Wöhrle S, et al. FGFR Genetic Alterations Predict for Sensitivity to NVP-BGJ398, a Selective Pan-FGFR Inhibitor. Cancer Discovery. 2012;2(12):1118–1133.

20. Lassman AB, Sepúlveda-Sánchez JM, Cloughesy TF, et al. Infigratinib in Patients with Recurrent Gliomas and FGFR Alterations: A Multicenter Phase II Study. Clin Cancer Res. 2022;28(11):2270–2277.

21. Javle M, Roychowdhury S, Kelley RK, et al. Infigratinib (BGJ398) in previously treated patients with advanced or metastatic cholangiocarcinoma with FGFR2 fusions or rearrangements: mature results from a multicentre, open-label, single-arm, phase 2 study. Lancet Gastroenterol Hepatol. 2021;6(10):803–815.

22. Abou-Alfa GK, Borbath I, Roychowdhury S, et al. PROOF 301: Results of an early discontinued randomized phase 3 trial of the oral FGFR inhibitor infigratinib vs. gemcitabine plus cisplatin in patients with advanced cholangiocarcinoma (CCA) with an FGFR2 gene fusion/rearrangement. J Clin Oncol. 2024;42(3_suppl):516–516.

23. Pal SK, Grivas P, Gupta S, et al. Infigratinib versus placebo in patients with resected urothelial cancer (UC) bearing FGFR3 mutation or fusion: Primary DFS analysis from the phase 3, randomized PROOF302 study. J Clin Oncol. 2024;42(4_suppl):629–629.

24. Hoyer-Kuhn H. Oral Infigratinib in Children with Achondroplasia - Targeted Treatment. N Engl J Med. 2025;392(9):920–922.

25. Gaudio E, Tarantelli C, Spriano F, et al. Targeting CD205 with the antibody drug conjugate MEN1309/OBT076 is an active new therapeutic strategy in lymphoma models. Haematologica. 2020;105(11):2584–2591.

26. Sartori G, Tarantelli C, Spriano F, et al. The ATR inhibitor elimusertib exhibits anti-lymphoma activity and synergizes with the PI3K inhibitor copanlisib. Br J Haematol. 2024;204(1):191–205.

27. Andrews S. FastQC A Quality Control tool for High Throughput Sequence Data; 2010.

28. Bolger AM, Lohse M, Usadel B. Trimmomatic: a flexible trimmer for Illumina sequence data. Bioinformatics. 2014;30(15):2114–2120.

29. Dobin A, Davis CA, Schlesinger F, et al. STAR: ultrafast universal RNA-seq aligner. Bioinformatics. 2013;29(1):15–21.

30. Anders S, Pyl PT, Huber W. HTSeq--a Python framework to work with high-throughput sequencing data. Bioinformatics. 2015;31(2):166–169.

31. Robinson MD, McCarthy DJ, Smyth GK. edgeR: a Bioconductor package for differential expression analysis of digital gene expression data. Bioinformatics. 2010;26(1):139–140.

32. Law CW, Chen Y, Shi W, Smyth GK. voom: Precision weights unlock linear model analysis tools for RNA-seq read counts. Genome Biol. 2014;15(2):R29.

33. Subramanian A, Tamayo P, Mootha VK, et al. Gene set enrichment analysis: a knowledge-based approach for interpreting genome-wide expression profiles. Proc Natl Acad Sci U S A. 2005;102(43):15545–15550.

34. Shaffer AL, Wright G, Yang L, et al. A library of gene expression signatures to illuminate normal and pathological lymphoid biology. Immunol Rev. 2006;210:67–85.

35. Johnson Z, Tarantelli C, Civanelli E, et al. IOA-244 is a Non-ATP-competitive, Highly Selective, Tolerable PI3K Delta Inhibitor That Targets Solid Tumors and Breaks Immune Tolerance. Cancer Res Commun. 2023;3(4):576–591.

36. Young RM, Staudt LM. Targeting pathological B cell receptor signalling in lymphoid malignancies. Nat Rev Drug Discov. 2013;12(3):229–243.

37. Arribas AJ, Napoli S, Cascione L, et al. ERBB4-Mediated Signaling Is a Mediator of Resistance to PI3K and BTK Inhibitors in B-cell Lymphoid Neoplasms. Mol Cancer Ther. 2024;23(3):368–380.

38. Arribas A, Napoli S, Cascione L, et al. Secondary resistance to the PI3K inhibitor copanlisib in marginal zone lymphoma. European Journal of Cancer. 2020;138:S40–S40.

39. Arribas AJ, Guidetti F, Cannas E, et al. IL-16 production is a mechanism of resistance to BTK inhibitors and R-CHOP in lymphomas. bioRxiv. 2025.

40. Arribas AJ, Napoli S, Cascione L, et al. Resistance to PI3Kdelta inhibitors in marginal zone lymphoma can be reverted by targeting the IL-6/PDGFRA axis. Haematologica. 2022;107(11):2685–2697.

41. Knox C, Wilson M, Klinger CM, et al. DrugBank 6.0: the DrugBank Knowledgebase for 2024. Nucleic Acids Res. 2024;52(D1):D1265–d1275.

42. Musa A, Tripathi S, Dehmer M, Emmert-Streib F. L1000 Viewer: A Search Engine and Web Interface for the LINCS Data Repository. Front Genet. 2019;10:557.

43. Yang W, Soares J, Greninger P, et al. Genomics of Drug Sensitivity in Cancer (GDSC): a resource for therapeutic biomarker discovery in cancer cells. Nucleic Acids Res. 2013;41(Database issue):D955–961.

