## Supplementary figures for "Targeting Aberrant FGFR Signaling with Infigratinib Enhances the Efficacy of BTK/PI3K Inhibitors and Bendamustine in Lymphoma Cells"

**Supplementary Table 1. IC50.**
**Supplementary Table 2. Combination data.**
**Supplementary Table 3. Limma and GSEA.**

**Supplementary Figure 1. FGFR expression in lymphoma cell lines: A)** Heatmap showing expression of FGFRs and FGFs across a pannel of cell lines derived from MCL, ABC-DLBCL and germinal center B-cell (GCB) DLBCL **B)** Immunoblot for FGFR1, FGFR2, FGFR3 and FGFR4 protein in a subset of selected cell lines and **C)** its protein quantification expressed as ratio to SP53.

A) FGFRs and FGFs in all cell lines

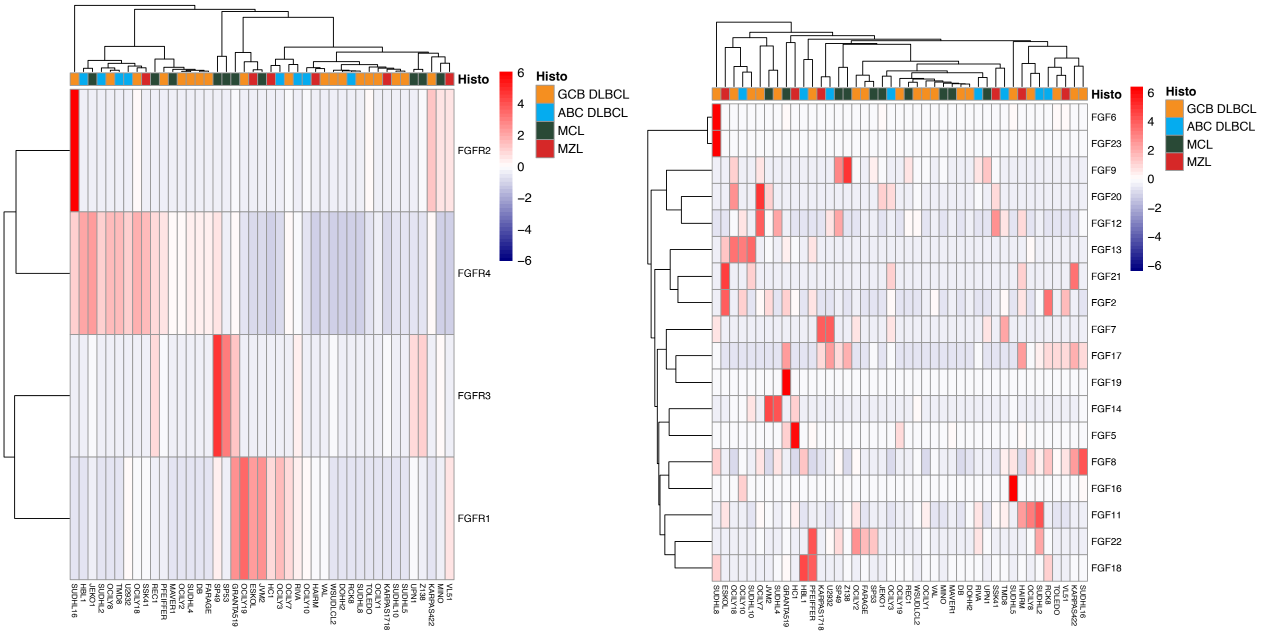


B

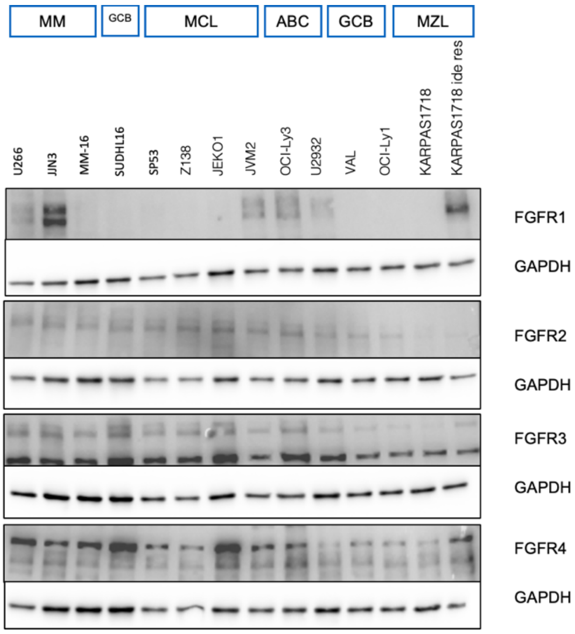

C)

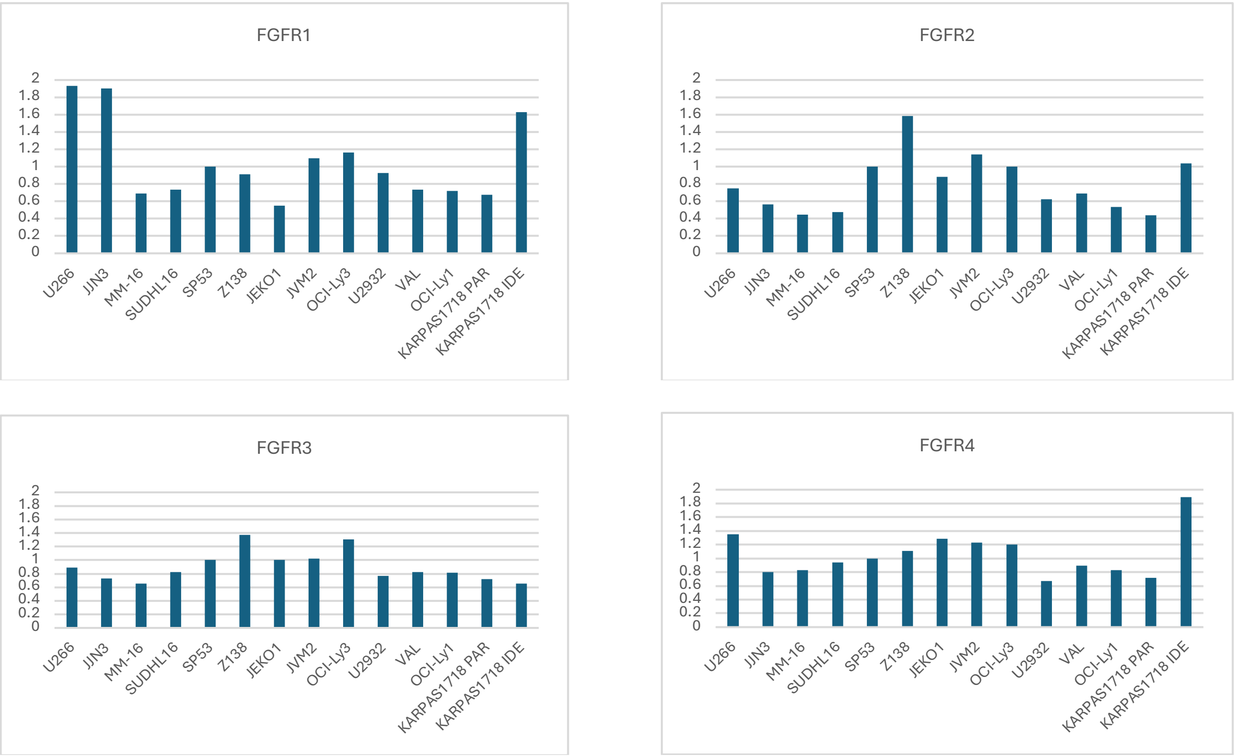
FGFR1
2
1.8
1.6
1.4
1.2
1
0.8
0.6
0.4
0.2
0
U266
JJN3
MM-16
SUDHL16
SP53
Z138
JEKO1
JVM2
OCI-Ly3
U2932
VAL
OCI-Ly1
KARPAS1718 PAR
KARPAS1718 IDE
FGFR2
2
1.8
1.6
1.4
1.2
1
0.8
0.6
0.4
0.2
0
U266
JJN3
MM-16
SUDHL16
SP53
Z138
JEKO1
JVM2
OCI-Ly3
U2932
VAL
OCI-Ly1
KARPAS1718 PAR
KARPAS1718 IDE
FGFR3
2
1.8
1.6
1.4
1.2
1
0.8
0.6
0.4
0.2
0
U266
JJN3
MM-16
SUDHL16
SP53
Z138
JEKO1
JVM2
OCI-Ly3
U2932
VAL
OCI-Ly1
KARPAS1718 PAR
KARPAS1718 IDE
FGFR4
2
1.8
1.6
1.4
1.2
1
0.8
0.6
0.4
0.2
0
U266
JJN3
MM-16
SUDHL16
SP53
Z138
JEKO1
JVM2
OCI-Ly3
U2932
VAL
OCI-Ly1
KARPAS1718 PAR
KARPAS1718 IDE

**Supplementary Figure 2. Single-agent activity of infigratinib in lymphoma cell lines: A)** Drug response curves of infigratinib in MCL **B)** ABC and GCB DLBCL models, and **C)** MZL cell lines, including **D)** KARPASS178 and **E)** VL51 with acquired resistance to PI3K, BTK or PI3K/BCL2 inhibitors. Viability was assessed by MTT assay upon 72 hours of exposure to increasing doses of infigratinib. Curves correspond to the mean of at least two independent experiments. Error bars represent the standard deviation of the mean. Supp Table 1 shows the IC50 values of infigratinib upon 72 hours of exposure (4-parameter log-regression equation, "drc" package in R environment).

A)

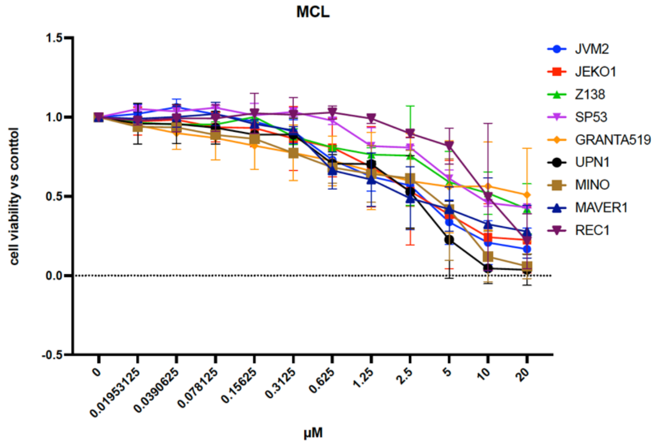


B)

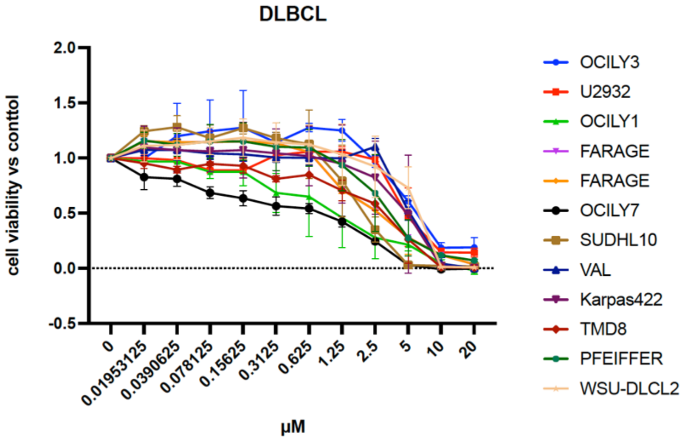


C)

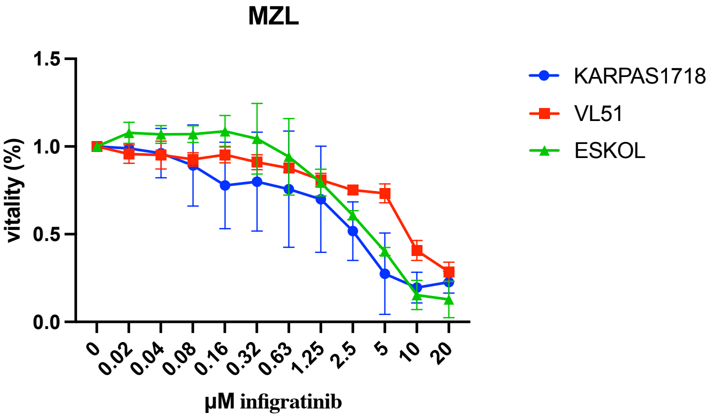

D)

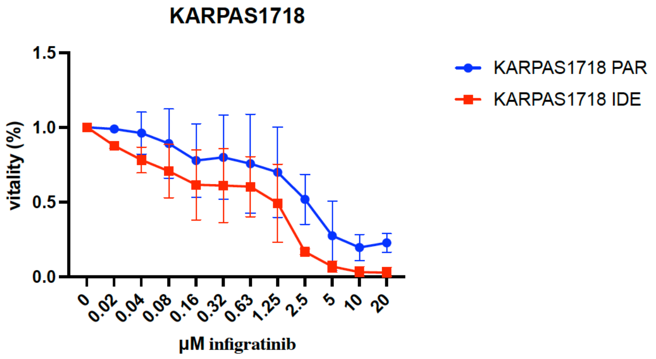
KARPAS1718
1.5
1.0
0.5
0.0
vitality (%)
0
0.02
0.04
0.08
0.16
0.32
0.63
1.25
2.5
5
10
20
µM infigratinib
KARPAS1718 PAR
KARPAS1718 IDE

E)

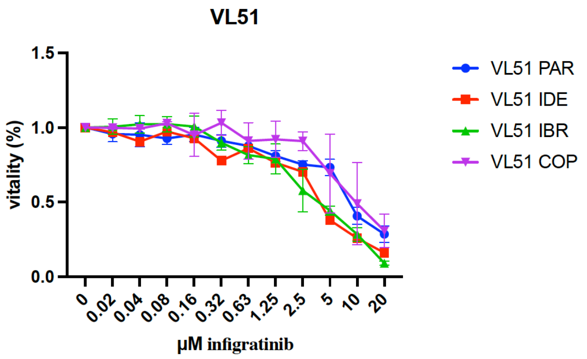
VL51
1.5
1.0
0.5
0.0
vitality (%)
0
0.02
0.04
0.08
0.16
0.32
0.63
1.25
2.5
5
10
20
µM infigratinib
VL51 PAR
VL51 IDE
VL51 IBR
VL51 COP

**Supplementary Figure 3. Infigratinib leads to decreased percentage of proliferating lymphoma cells.** Cell cycle phases upon infigratinib assessed by PI staining (FACS) obtained in four GCB (OCI-LY-7, PFEIFFER, FARAGE and VAL), two ABC DLBCL (TMD8 and U2932) in addition to Karpas1718 and its derivative with acquired resistance to BTK and PI3K inhibitors.

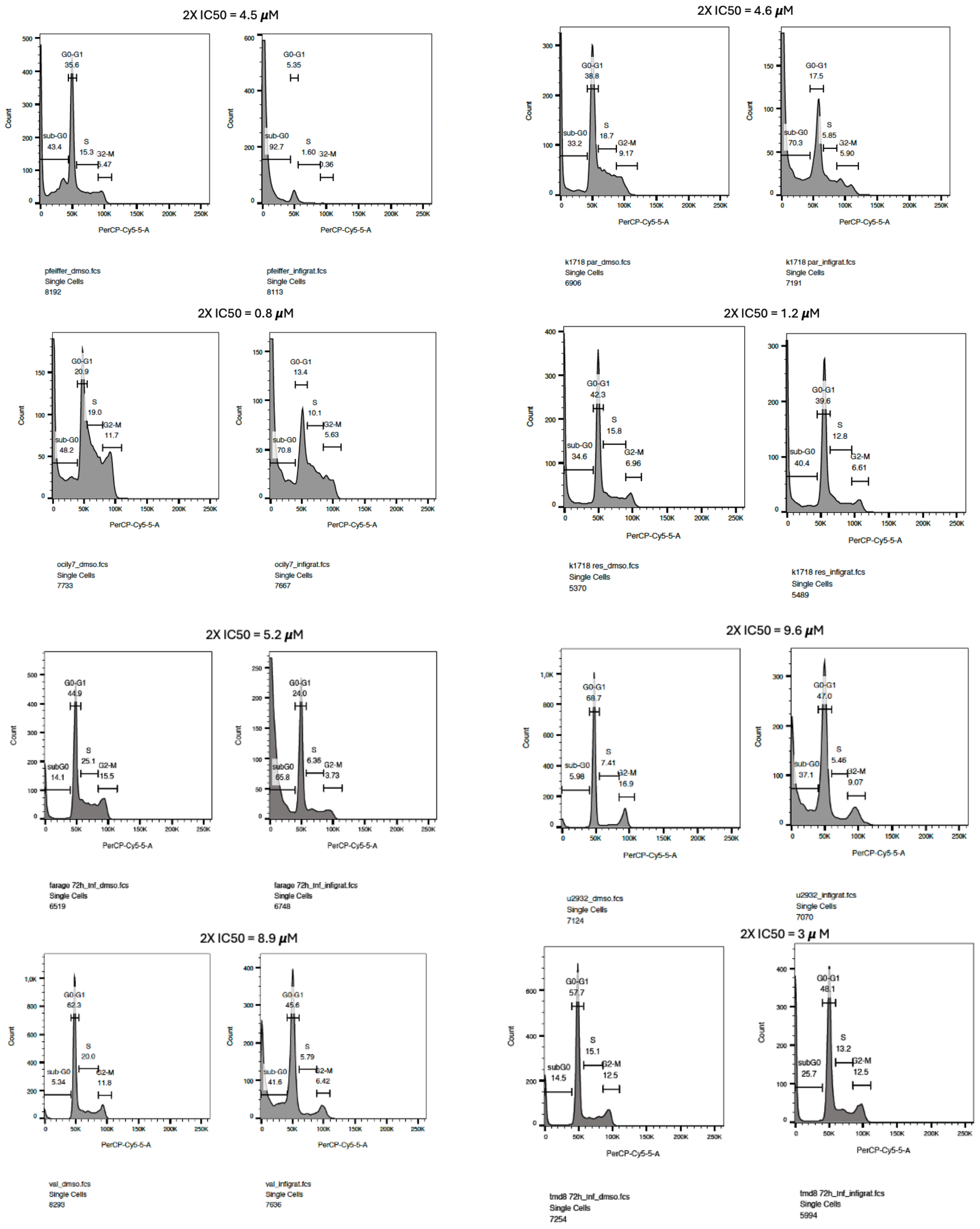

**Supplementary Figure 4. Ibrutinib and infigratinib are targeting two complementary pathways, the BCR signalling and the cell cycle pathway respectively.** Differentially enriched hallmark genesets (MSigDB, Broad Institute) across treatments. FDR for false discovery rate, NES for normalized enrichment score.

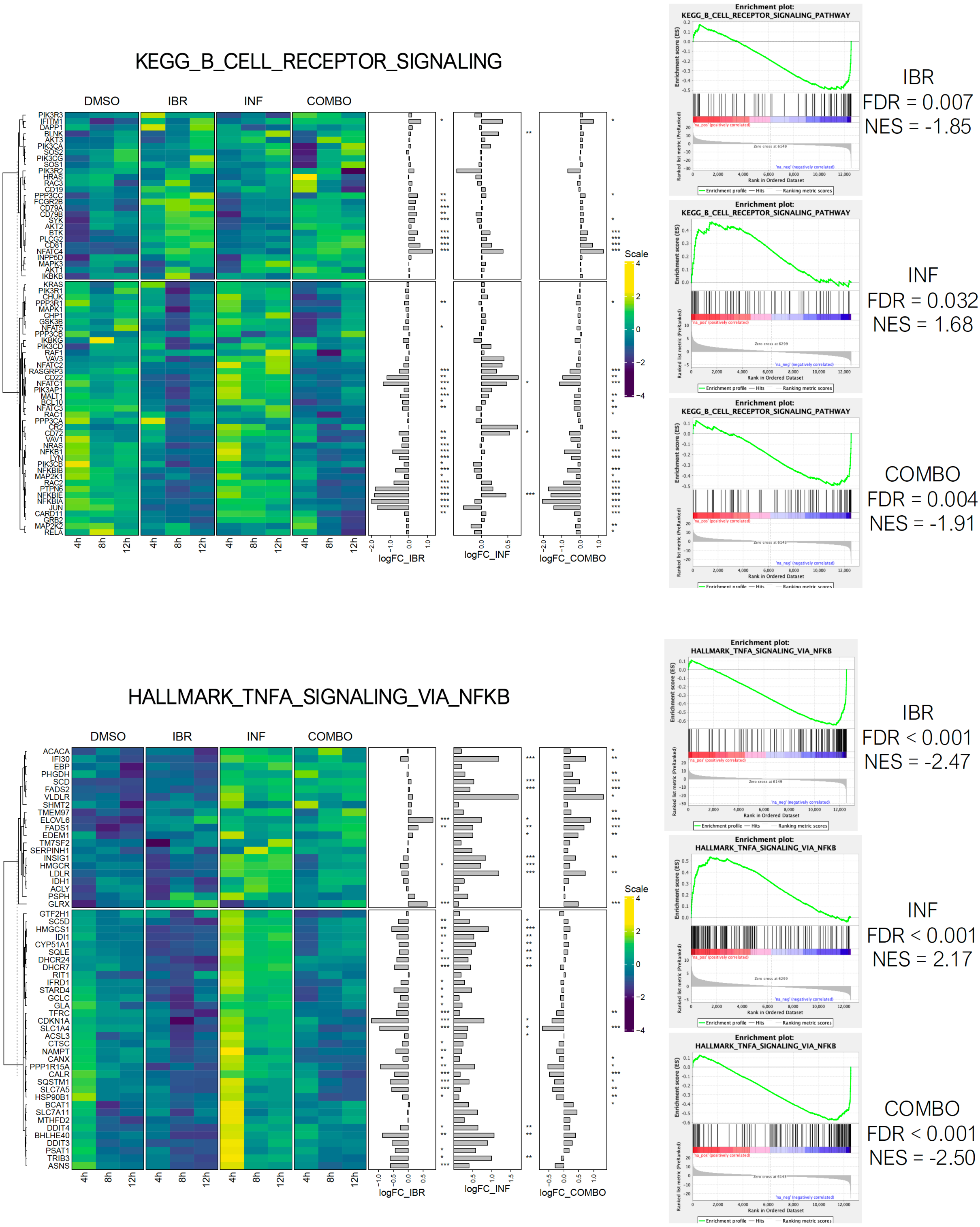

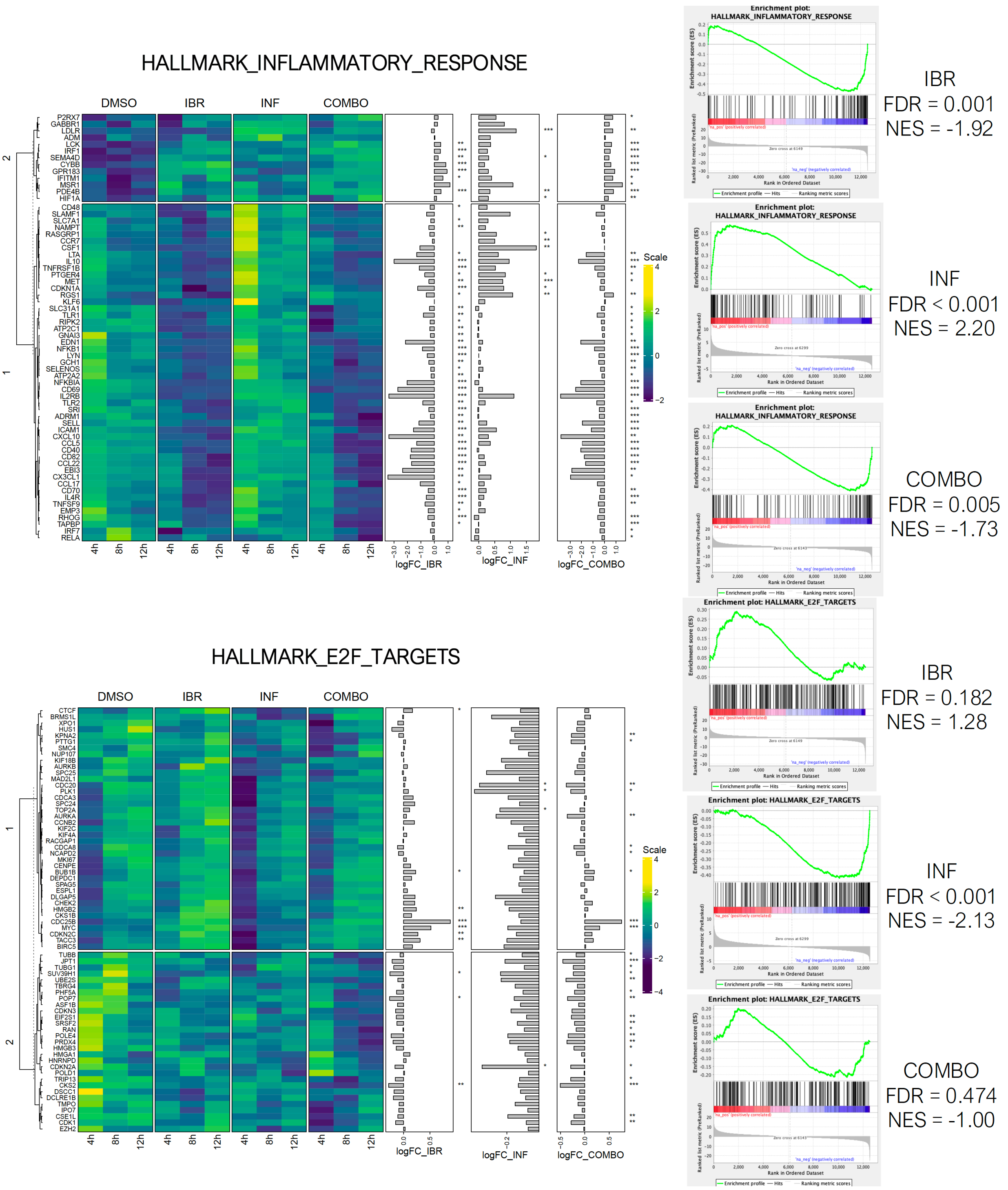
HALLMARK_INFLAMMATORY_RESPONSE
DMSO
IBR
INF
COMBO
4h
8h
12h
logFC_IBR
logFC_INF
logFC_COMBO
Scale
Enrichment plot: HALLMARK_INFLAMMATORY_RESPONSE
IBR
FDR = 0.001
NES = -1.92
INF
FDR < 0.001
NES = 2.20
COMBO
FDR = 0.005
NES = -1.73
HALLMARK_E2F_TARGETS
Enrichment plot: HALLMARK_E2F_TARGETS
IBR
FDR = 0.182
NES = 1.28
INF
FDR < 0.001
NES = -2.13
COMBO
FDR = 0.474
NES = -1.00

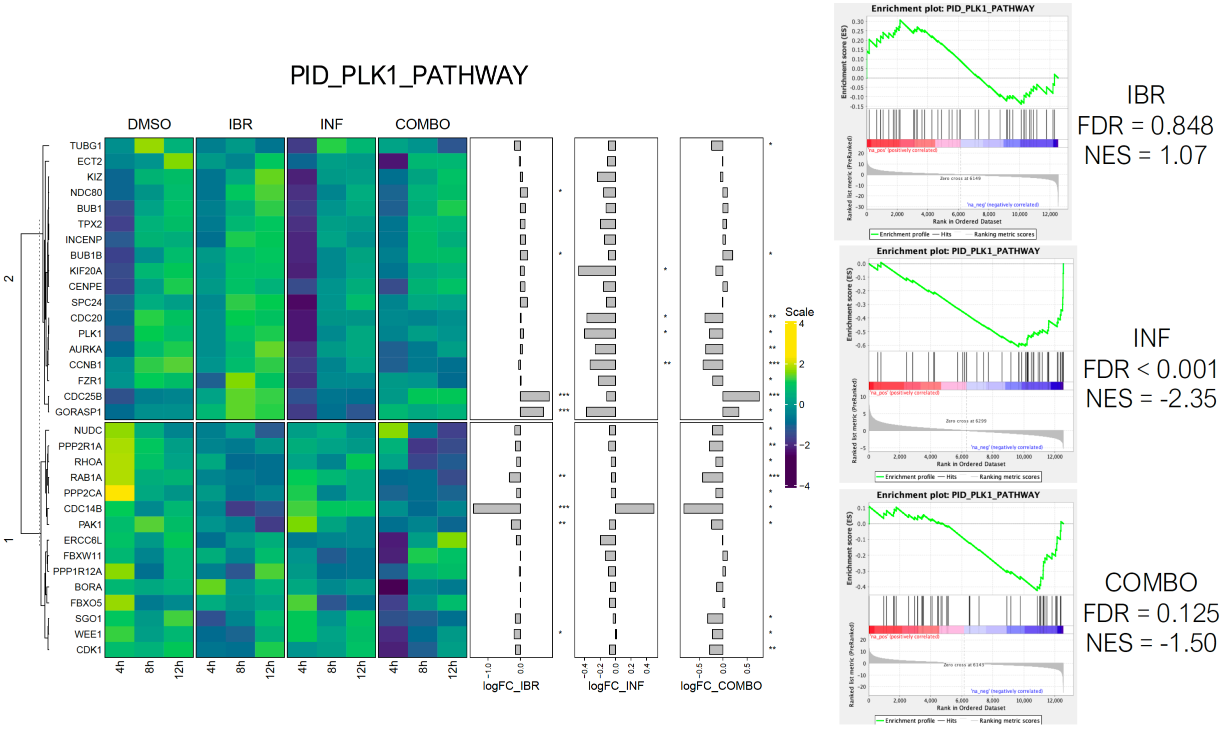
PID_PLK1_PATHWAY
DMSO
IBR
INF
COMBO
TUBG1
ECT2
KIZ
NDC80
BUB1
TPX2
INCENP
BUB1B
KIF20A
CENPE
SPC24
CDC20
PLK1
AURKA
CCNB1
FZR1
CDC25B
GORASP1
NUDC
PPP2R1A
RHOA
RAB1A
PPP2CA
CDC14B
PAK1
ERCC6L
FBXW11
PPP1R12A
BORA
FBXO5
SGO1
WEE1
CDK1
4h
8h
12h
logFC_IBR
logFC_INF
logFC_COMBO
Scale
Enrichment plot: PID_PLK1_PATHWAY
IBR
FDR = 0.848
NES = 1.07
INF
FDR < 0.001
NES = -2.35
COMBO
FDR = 0.125
NES = -1.50

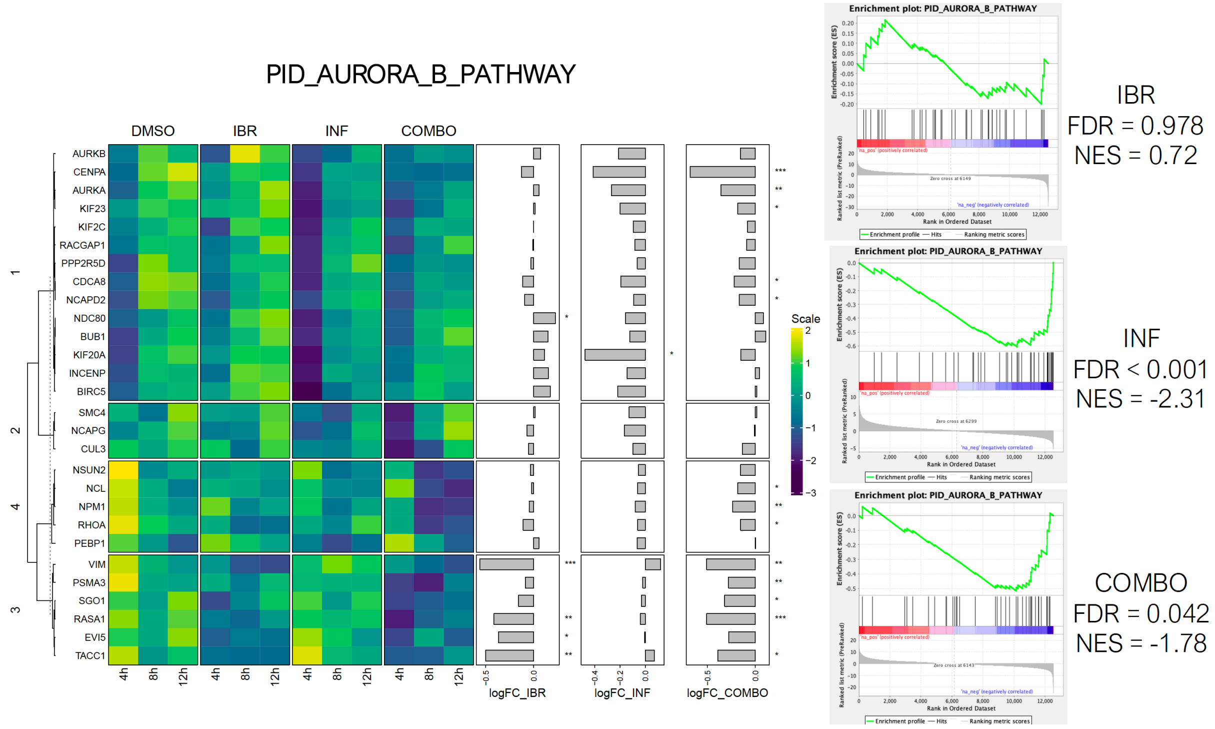
PID_AURORA_B_PATHWAY
DMSO
IBR
INF
COMBO
AURKB
CENPA
AURKA
KIF23
KIF2C
RACGAP1
PPP2R5D
CDCA8
NCAPD2
NDC80
BUB1
KIF20A
INCENP
BIRC5
SMC4
NCAPG
CUL3
NSUN2
NCL
NPM1
RHOA
PEBP1
VIM
PSMA3
SGO1
RASA1
EVI5
TACC1
4h
8h
12h
logFC_IBR
logFC_INF
logFC_COMBO
Scale
Enrichment plot: PID_AURORA_B_PATHWAY
IBR
FDR = 0.978
NES = 0.72
INF
FDR < 0.001
NES = -2.31
COMBO
FDR = 0.042
NES = -1.78

**Supplementary Figure 5. Infigratinib and ibrutinib combination determines transcriptome changes underscoring complementary pathway inhibition.** Transcriptional changes induced by infigratinib treatment were compared to those induced by other drugs present in publicly available data, such as the LINCS L1000 project (L1000) and the Genomics of Drug Sensitivity in Cancer (GDSC) project. PP-2 is an inhibitor for Src-family kinases; PI-103 is a dual PI3K and mTOR inhibitor.

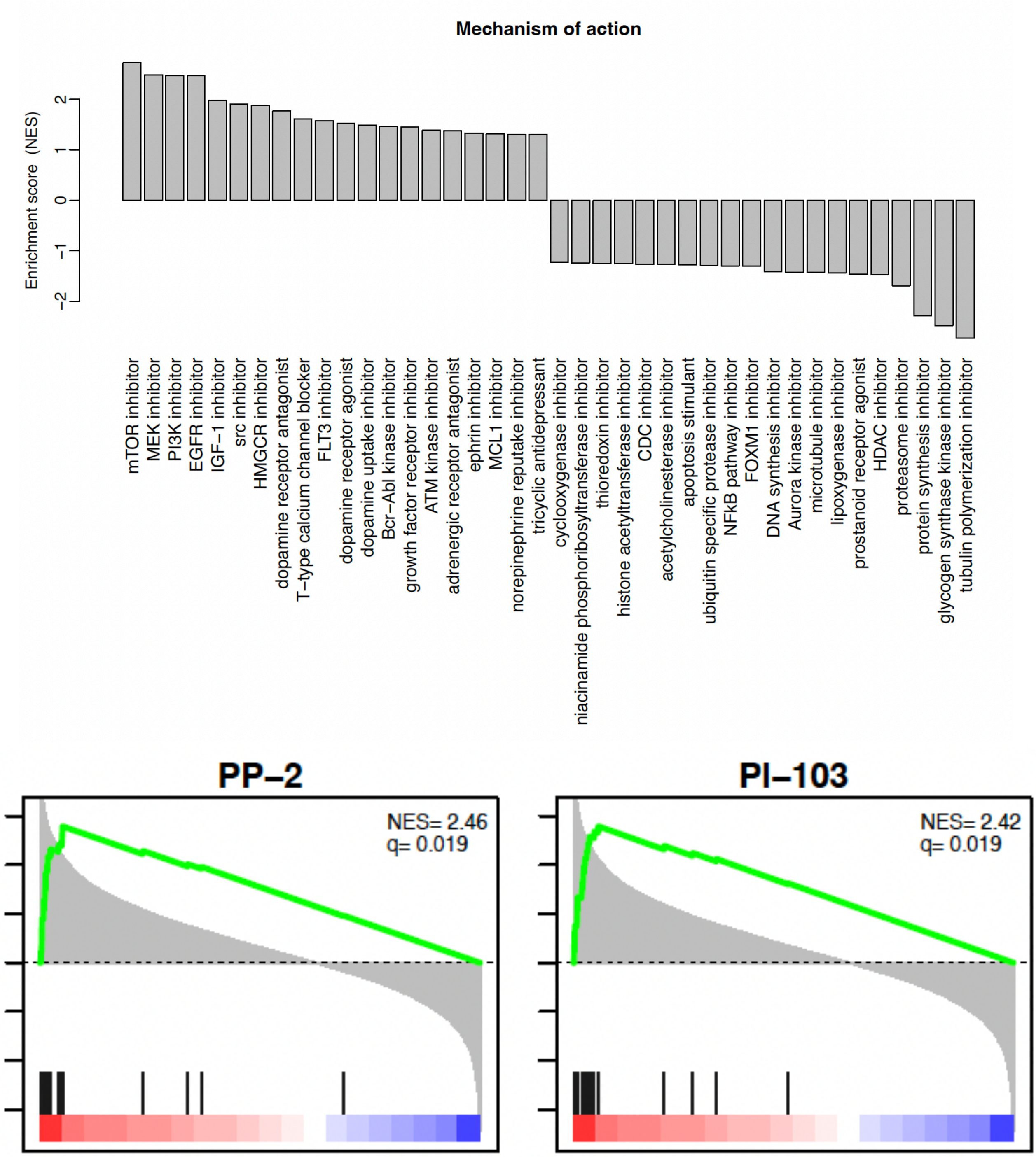
